# Temporal-deviation-driven community detection uncovers early-warning signals for critical transitions in complex diseases

**DOI:** 10.64898/2026.05.08.723925

**Authors:** Letian Wang, Minrui Xu, Hao Yan, Yan Zheng, Shuang Feng, Yiming Zhang, Chang Li, Dehui Qiu, Bin Hu, Xiaohua Wan, Fa Zhang

**Author notes:** Contributing authors.

## Abstract

Early detection of critical transitions in complex diseases is crucial for timely clinical intervention. However, as patients often provide only a single snapshot, identifying sample-specific early-warning signals (EWS) from a dynamical evolution perspective remains challenging, coupled with high-dimensional noise amplification. Here, we present TD-COM, a framework for detecting personalized EWS of critical transitions via single-sample community detection. By constructing a temporal perturbation map STDN, TD-COM captures latent dynamical perturbations inferred from static individual profiles. Synergizing these temporal-deviation signals with static topological features, TD-COM implements a multilevel node filtering strategy during community detection, effectively suppressing single-sample noise. Validated on hour-scale, multi-year, and multi-decade transcriptomic data, TD-COM robustly detects critical states preceding clinical deterioration and uncovers their underlying molecular mechanisms. Comparative experiments demonstrate that TD-COM outperforms existing methods in accuracy and topological robustness. Thus, TD-COM provides a generalizable framework for personalized early warning of complex diseases, particularly when longitudinal sampling is infeasible.

## 1 Introduction

There is growing recognition that many natural processes do not evolve linearly or gradually. Instead, they often undergo abrupt, qualitative shifts at critical thresholds, a phenomenon known as a “critical transition” ^1–3^, manifesting across time scales. Non-linear dynamical models of disease progression reveal three distinct states ^4,5^ (Figure 1a): a relatively healthy state, a critical state, and a deteriorated state. The healthy and deteriorated states are stable, while the critical state is not. Poised at the edge of stability, the critical state is characterized by critically elevated sensitivity to perturbations, thereby predisposing the system to irreversible collapse. Recognizing this critical state and detecting patient-specific early-warning signals (EWS) not only provide a window for preemptive intervention, but also enable the stratification of critical-state subtypes and the identification of shared molecular drivers, insights that can guide both trajectory-specific and broad-spectrum therapeutic strategies.

**Fig. 1.**
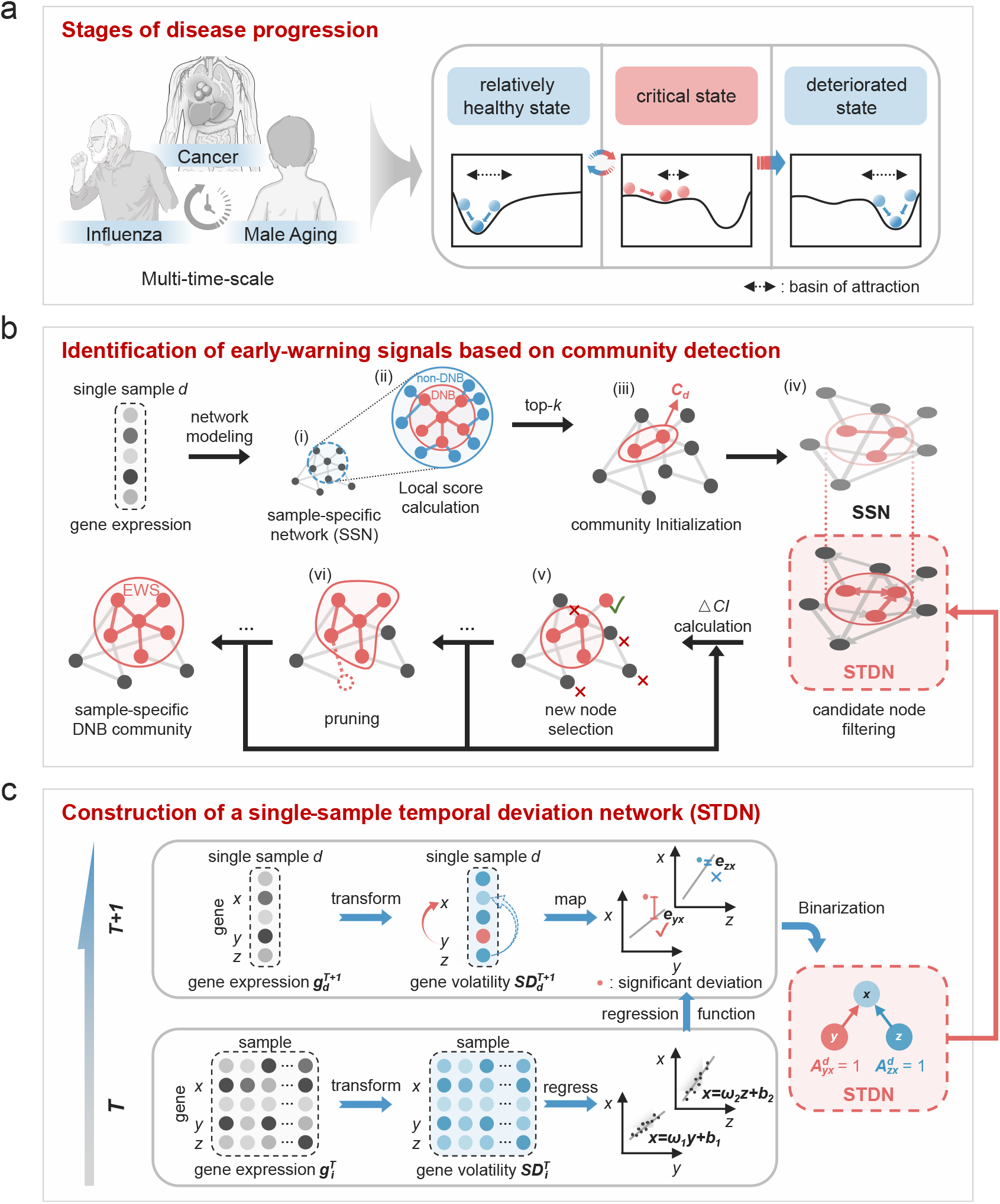
Overview of TD-COM. **a**. The progression of complex diseases across diverse timescales can be modeled as a nonlinear dynamical system, typically undergoing three stages: relatively healthy, critical, and deteriorated states. The critical state is highly sensitive to perturbations; once this tipping point is crossed, the system often undergoes an abrupt and irreversible catastrophic shift. Therefore, identifying early-warning signals (EWS) preceding such critical transitions in individual samples is essential for enabling personalized early intervention. **b**. Based on the sample-specific network (SSN) constructed from single-sample expression data (Figure 1b(i)), each gene is first scored according to its topological significance within the local neighborhood (Figure 1b(ii)). High-scoring nodes are subsequently selected as initial seeds to initiate the DNB community *C*_*d*_ (Figure 1b(iii)). To mitigate error propagation during community detection, a multi-level node filtering strategy is implemented, encompassing candidate node screening (Figure 1b(iv)), dynamic threshold optimization (Figure 1b(v)), and iterative node pruning (Figure 1b(vi)). The resulting DNBs within the refined *C*_*d*_ constitute the sample-specific EWS for the disease. **c**. We propose a single-sample temporal deviation network (STDN), which leverages population-level temporal prior information from stage *T* to assess, within a single sample at stage *T* + 1, whether a target gene exhibits a significant nonlinear deviation from the expected gene fluctuation pattern established at stage *T* . By treating adjacent disease stages as dynamic references, STDN captures latent temporal dynamics in a single snapshot, enabling sample-specific EWS extraction across diseases with diverse progression timescales.

Identifying patient-specific EWS of critical transitions remains a formidable challenge. As disease progression is inherently dynamic, effective warning methods must capture dynamical instabilities rather than static snapshots. Yet most clinical indicators merely contrast healthy and diseased states, reflecting the outcome of a transition rather than its dynamic precursors ^6^. This challenge is compounded by data limitations: traditional EWS identification requires longitudinal sampling from individual patients ^7^, which is often prohibitive due to cost and poor compliance. Most patients contribute only a single biosample. Consequently, discerning dynamical instabilities embedded in isolated samples to enable personalized EWS inference remains difficult.

In this context, methods such as SSN^8^ and SWEET^9^ reconstruct sample-specific gene regulatory networks from static snapshots. Community detection algorithms have proven effective in analyzing such networks ^10,11^, showing potential for single-sample EWS. This potential is grounded in extensive evidence that tipping events in complex networks originate in highly cohesive subnetworks. Such synchronous destabilization then cascades through network coupling, ultimately driving global transitions ^12–16^. However, current community detection methods rely primarily on static topological measures such as modularity^17,18^, thus neglecting the dynamical features inherent to critical transitions. Consequently, these methods often fail to identify precursor communities driving state shifts, thereby obscuring disease-dynamical signatures of sample-specific EWS.

In recent years, the dynamical network biomarker theory (DNB, see Supplementary Section IX.d)^6^, grounded in nonlinear dynamics, has proven effective in capturing pre-transition states from transcriptomic data. Notably, transcriptomic data capture genome-wide expression dynamics, offering a sensitive, system-wide perspective on impending transitions ^19^. The applicability of DNB theory spans diverse biological contexts, including cell fate decisions ^20,21^, COVID-19^22^, and various cancers ^23–25^. Building on this theory, Wang et al. proposed the com-DNB and crsDNB methods ^26,27^ to incorporate dynamical constraints into community detection for single-sample networks. Yet these methods face an inherent limitation: sample-specific EWS should ideally be inferred from the temporal evolution of a system approaching critical transition, thereby capturing coordinated variability over time. Instead, existing approaches rely solely on information derived from static snapshots, yielding characterizations that fail to capture the true destabilization dynamics. Moreover, noise in high-dimensional transcriptomic data is amplified under single-sample limitations, causing false-positive nodes to be assigned to communities and triggering error cascades. Therefore, a robust framework leveraging genuine dynamical properties is essential for reliably extracting sample-specific EWS.

Here, we present TD-COM (temporal-deviation-driven community-detection), a framework that accurately identifies critical states in complex diseases and robustly detects personalized EWS from single-sample transcriptomic data. Unlike conventional methods, TD-COM captures latent dynamic perturbation signatures within single-sample data, thereby identifying critical-state biomarker communities while effectively suppressing noise accumulation inherent in such data. Specifically, TD-COM constructs a Single-sample Temporal Deviation Network (STDN), a temporal perturbation map at the single-sample level, to characterize the temporal variations of system variables inferred from static observations. By synergizing this temporal information with static topological characteristics, TD-COM implements a multi-level node filtering strategy during community detection, enabling noise-suppressed identification of sample-specific EWS that reflect authentic dynamic critical behaviors.

We applied TD-COM to seven real transcriptomic datasets spanning hour-scale influenza virus infection, multi-year cancer progression, and multi-decade male aging. In influenza infection, TD-COM detected sample-specific EWSs at least 16 hours before symptom onset, enabling preemptive antiviral intervention. Across five cancer types with multi-year progression, TD-COM identified pre-deterioration critical states. Specifically, we uncovered two immune-fibrotic critical-state subtypes in LUSC and shared drivers of malignant transition between LUSC and LUAD, guiding both personalized and broad-spectrum therapeutic strategies. In a male aging cohort, TD-COM identified critical windows at ages 61–65 and 71–75 in PBMC and classical monocyte transcriptomes, revealing core pathways underlying age-related decline and offering precise temporal targets for delaying aging. Compared with existing methods and ablation studies, TD-COM more effectively characterizes system instability at critical states with high computational robustness and efficiency. Overall, TD-COM reliably captures critical transitions across multiple timescales and serves as a powerful early-warning tool.

## 2 Results

### 2.1 Overview of the TD-COM

We present TD-COM (temporal-deviation-driven community-detection) to enable robust detection of patient-specific EWS from a truly dynamic perspective using single-sample transcriptomic data. As illustrated in Figure 1b, TD-COM employs a multi-stage DNB community detection pipeline, integrating the dynamics-informed STDN (Single-sample Temporal Deviation Network) with the conventional static network SSN to identify sample-specific DNB communities: first, communities are initialized around high-scoring seed nodes (Figure 1b(ii)-(iii)); next, the multi-level node filtering strategy is applied iteratively through three steps: (1) dual filtering of candidate nodes based on complementary information from the STDN and SSN (Figure 1b(iv)); (2) adaptive dynamic gain thresholding (Figure 1b(v)); and (3) pruning operations (Figure 1b(vi)). By synergistically integrating the three filtering steps, TD-COM effectively suppresses noise inherent in single-sample data and mitigates error propagation, thereby enabling reliable identification of single-sample DNB community that herald system instability preceding the critical transition. Recurrent nodes within single-sample DNBs of critical-state samples are designated as population-DNBs.

The core innovation of TD-COM lies in the construction of the STDN, which extracts latent dynamical information from static single-sample data (Figure 1c). The core innovation of TD-COM lies in the construction of the STDN, which extracts latent dynamical information from static single-sample data (Figure 1c). For an individual sample at stage *T* +1, the STDN leverages population-level time-series prior knowledge from stage *T* to quantify the extent to which each gene’s expression volatility deviates from the expected pattern established at stage *T* . By using the adjacent stage as a dynamic reference, this approach reveals latent, nonlinear dynamical perturbations that herald an impending critical transition across complex disease processes operating at multiple timescales. These perturbations serve as dynamical signatures of variable instability, enabling reliable detection of an approaching critical transition even from a single transcriptomic snapshot. Thus, TD-COM delineates sample-specific EWS in the context of disease progression. These perturbations serve as dynamical signatures of variable instability, enabling reliable detection of an approaching critical transition even from a single transcriptomic snapshot. Thus, TD-COM delineates sample-specific EWS in the context of disease progression.

### 2.2 Analysis of multi-year cancer progression using TD-COM

#### 2.2.1 Identification of critical states in cancer progression

As shown in Figure 2, we applied TD-COM to gene expression data from five cancer types in the TCGA database. Normal samples were served as the reference set, while tumor samples were stratified by TNM stage for LUSC, LUAD, READ, and COAD, and by FIGO stage for UCEC (see Supplementary Table 1). TD-COM identified a personalized gene community and its corresponding *CI* for each tumor sample. Subsequently, statistical significance testing was employed to detect population-level critical states. Specifically, TD-COM pinpointed stage IB in LUSC, stage IIB in LUAD, stage IIIC in UCEC, stage II in READ, and stage IIA in COAD as the respective critical states (Figure 2a-e).

**Fig. 2.**
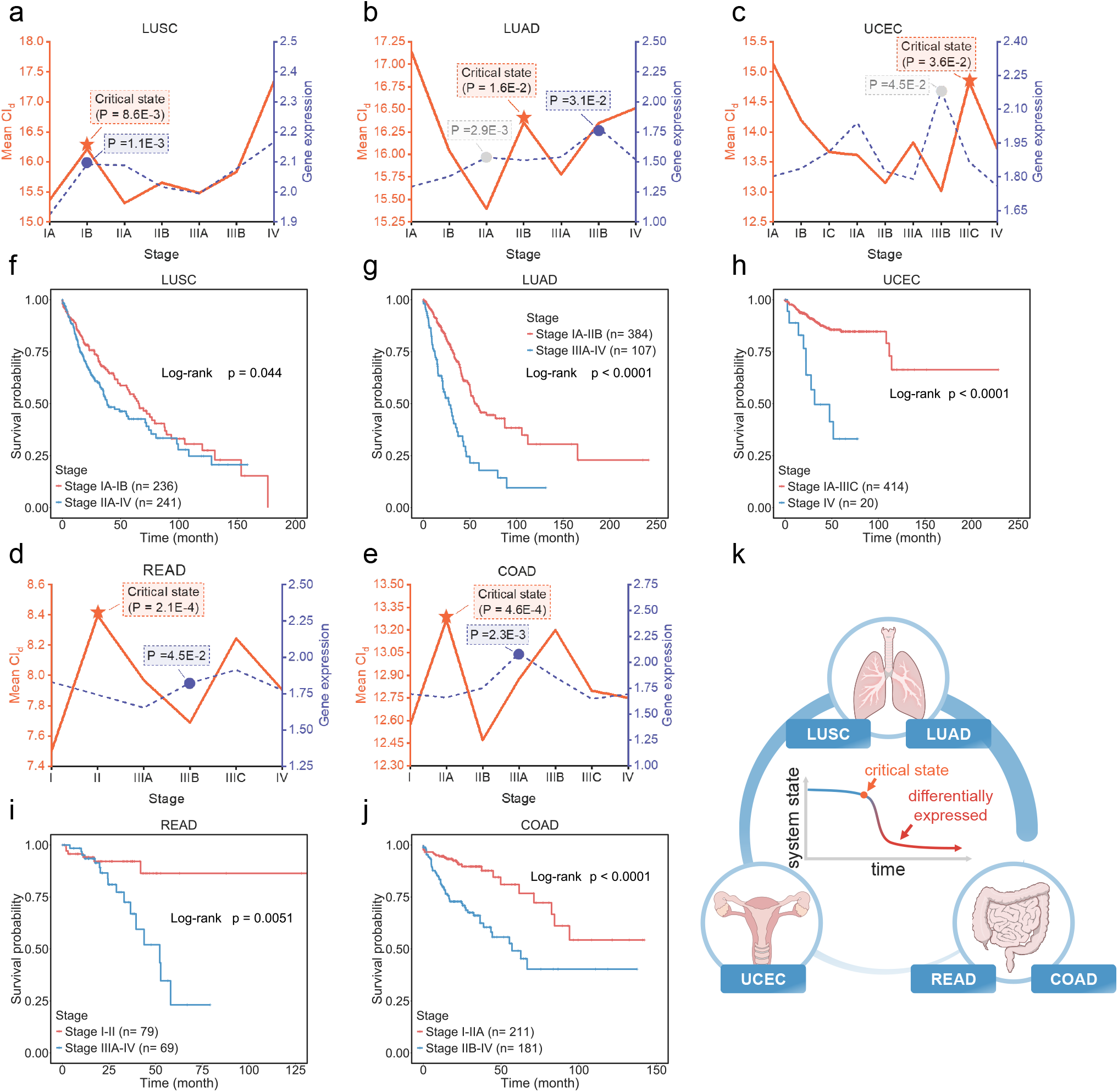
TD-COM predicts impending critical transitions in multi-year cancer progressions. TD-COM successfully identified the critical states preceding catastrophic deterioration in **(a)** LUSC, **(b)** LUAD, **(c)** UCEC, **(d)** READ, and **(e)** COAD. The blue dots on the blue dashed lines represent the positions where significant changes in the expression of highly expressed genes occur, which typically lag behind the identified critical states. The gray dots correspond to the changes in *CI* prior to the critical states. Survival analyses in **(f)**–**(j)** reveal statistically significant differences in patient survival between the pre- and post-transition phases, thereby validating the EWSs detected by TD-COM. **(k)** TD-COM was applied to TCGA datasets for five common cancers from the respiratory, reproductive, and digestive systems.

As shown in Figure 2a, a critical shift occurred in LUSC following a sharp increase in the *CI* at stage IB (*P* = 8.6 × 10^*−*3^), suggesting that tumor cells were on the verge of breaching the primary tumor site and spreading to ipsilateral peribronchial or hilar lymph nodes for the first time ^28^. This transition marks a pivotal shift from surgery alone to multimodal therapy combining surgery with adjuvant systemic treatment. In LUAD, the *CI* increased sharply at stage IIB *P* = 1.6×10^*−*2^; Figure 2b), serving as an early-warning signal for subsequent metastasis to ipsilateral mediastinal or subcarinal lymph nodes at stage IIIA^28^. As illustrated in Figure 2c, when applied to UCEC, our method revealed a substantial increase in *CI* at stage IIIC (*P* = 3.6 × 10^*−*2^), after which tumors typically begin invading adjacent organs and spreading to distant sites ^29^. In READ, the *CI* exhibited a significant peak at stage II (*P* = 2.1 × 10^*−*4^), suggesting that the system approaches a tipping point at this stage (Figure 2d). As previously reported, regional lymph node metastasis emerges at stage IIIA in this cancer type ^30^. In COAD, the *CI* was significantly higher at stage IIA than at stage I (*P* = 4.6×10^*−*4^), indicating that stage IIA represents a critical state in disease progression (Figure 2e). Beyond this stage, the system undergoes an irreversible transition characterized by tumor invasion into the visceral peritoneum, which substantially increases the risk of intraperitoneal dissemination ^31^. In contrast, the blue dashed lines in Figures 2a–2e reflect the expression changes of differentially expressed genes (DEGs) during disease progression. Notably, the significant expression changes marked by blue dots lag behind the critical states shown as orange stars, making them unreliable early indicators of critical transitions. The gray dots marking significant changes at stage IIA (Figure 2b) and IIIB (Figure 2c) correspond to *CI* fluctuations that precede the identified critical states, rather than being driven by the critical states themselves. In UCEC, the increases at stages IIA and IIIB on the blue dashed lines correspond respectively to the increases at stages IA and IIIA on the orange lines. Additionally, Figure 2c lacks blue dots corresponding to the critical state at stage IIIC, as the disease has already progressed to an advanced phase where systemic collapse invalidates the traditional gene expression biomarkers that assume independent utility.

To validate the identified critical states, we performed survival analysis using the Kaplan-Meier method with the log-rank test. As shown in Figures 3f–3j, patients before the critical transition exhibited significantly longer survival times than those after it across all five cancer types (*P<*0.05). These results demonstrate that TD-COM is a robust method for detecting critical transitions and can deliver early, reliable warning signals prior to the irreversible progression of complex diseases. Further details are provided in Supplementary information III and XI.

## 3 Methods

### 3.1 Construction of a single-sample temporal deviation network

To uncover latent dynamical perturbations embedded in single-sample transcriptomic data, we constructed a Single-sample Temporal Deviation Network (STDN). The dataset is partitioned into a reference set (relatively healthy state) and multiple case sets (disease stages) arranged in chronological order.

For a target gene *x* and a reference gene *y* (Figure 1c), the expression volatility vectors across *m* samples at stage *T* are defined as 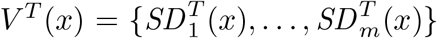 and 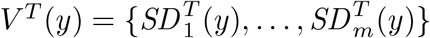, where *T* ≥ 0 (*T* = 0 indicates the reference set). 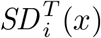 and 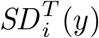 denote the volatility of genes *x* and *y* in sample *i* (1 ≤ *i* ≤ *m*).

For target gene *x*, gene *z* in Figure 1c is functionally equivalent to *y*, but we hereafter use the *x*–*y* pair as the representative example.

To assess the linear dependence between the fluctuation patterns of the two genes at stage *T*, we fitted a linear regression model with *V* ^*T*^ (*y*) as the explanatory variable and *V* ^*T*^ (*x*) as the response variable. Denoting 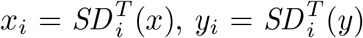 and *e*_*i*_ as the error term, we estimated the parameters using the least squares method (Equations (1)):

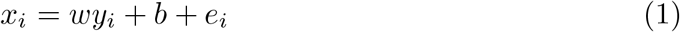

Subsequently, for sample *d* at stage *T* +1, the residual *e*_*yx*_ is defined as the difference between the actual volatility 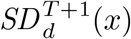 of target gene *x* and its model-predicted value. A significant deviation of *e*_*yx*_ from the baseline distribution constructed from stage *T* samples indicates that the volatility of target gene *x* in sample *d* cannot be effectively explained by the linear law established by reference gene *y* in the previous stage.

Thus, a directed STDN is constructed for sample *d*: a directed edge from reference gene *y* to target gene *x* is established when the volatility of *x* exhibits significant deviation relative to *y* (i.e., 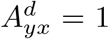 in adjacency matrix *A*^*d*^). Conversely, if no such deviation is observed relative to another reference gene *z*, no corresponding edge exists (i.e., 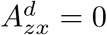 in Figure 1c). The STDN decodes latent disease evolutionary information from single-sample snapshots, thereby providing a dynamic perspective for candidate node filtering (Figure 1b(iv)). This filtering step is a critical component of the multilevel node filtering strategy in subsequent community detection based on the static SSN.

### 3.2 Identification of the DNB community in a single sample

As shown in Figure 1b, DNB community detection for sample *d* is performed within its SSN, guided by dynamic information from the previously constructed STDN. The construction of the SSN (Figure 1b(i)) is detailed in Supplementary Section VIII.e. This single-sample DNB community detection involves community initialization (Step 1, Figures 1b(ii–iii)) and a multi-level node filtering strategy (Steps 2–4, Figures 1b(iv)– (vi)), the latter of which effectively mitigates noise inherent in single-sample data. The DNBs within the final refined community constitute the sample-specific EWS for the disease.

#### [Step 1] Community initialization

In the SSN of sample *d*, we identify initial nodes by quantifying the likelihood of each gene *x* functioning as a DNB. Specifically, we construct a local subnetwork centered at each gene, comprising its first- and second-order neighbors. As shown in Figure 1b(ii), the center *x* and its first-order neighbors are heuristically designated as DNBs, while second-order neighbors are treated as non-DNBs. We then compute the local score 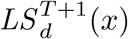 using Supplementary Equation (1), where *PCC*_*in*_ is restricted to edges between *x* and its first-order neighbors. As illustrated in Figure 1b(iii), we select the genes with the highest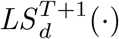 as the initial nodes of *C*_*d*_;

#### [Step 2] Candidate node filtering

We define the candidate set *S*_*d*_ as the first-order neighbors of *C*_*d*_ that are not yet community members. Each node *j* ∈ *S*_*d*_ is filtered from two complementary perspectives (Figure 1b(iv)): the static SSN topology and the STDN temporal dynamics, yielding two weights, *w*_SSN_(*j*) and *w*_STDN_(*j*) (Equations (2)–(3)). Here, *w*_SSN_(*j*) measures the mean connection strength between node *j* and community *C*_*d*_ in the SSN, whereas *w*_STDN_(*j*) quantifies the temporal deviation of *j* relative to *C*_*d*_ members serving as reference genes in the STDN. The *l* counts the *C*_*d*_ neighbors of *j* within the SSN, *α*(*i*) is the weight of node *i* ∈ *C*_*d*_, and *PCC*_*d*_(*i, j*) denotes the edge weight between nodes *i* and *j* in the SSN.

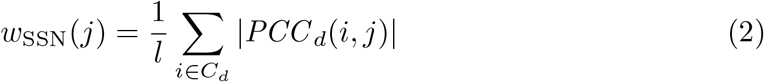

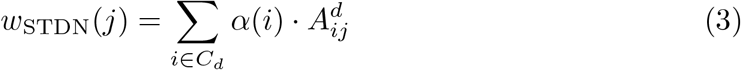

TD-COM employs a mode-aware filtering strategy based on *w*_SSN_ (Equation (4)), where *h* denotes the community size. The dense-mode threshold is prioritized to ensure robust connectivity, with the standard mode used only as a fallback when necessary. The threshold *θ*_STDN_ is set in a dataset-specific manner. Final candidate nodes must satisfy both *w*_SSN_ ≥ *θ*_SSN_ and *w*_STDN_ ≥ *θ*_STDN_. This dual-threshold mechanism filters for nodes that simultaneously satisfy two conditions: exhibiting strong coupling with existing *C*_*d*_ members in the SSN, and presenting critical dynamic signatures in the STDN that signal an impending transition.

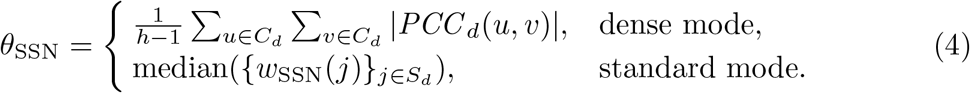

#### [Step 3] New node selection

TD-COM evaluates each candidate node by calculating the potential gain in *CI* from its inclusion, as defined in Equation (5). Here, intra-community volatility strength (*IVS* ), intra-community correlation strength (*ICS* ), and cross-community correlation strength (*CCS* ) correspond to the three characteristics of DNB theory, respectively, and *f* denotes the size of *S*_*d*_.

The minimum required gain threshold (*τ*_in_) for this round is then computed using Equation (6), where *τ*_0_ is the maximum gain factor. Here, *τ*_in_ is a dynamic gain threshold that adapts to the current community size *h*. Node selection then follows the operational mode:

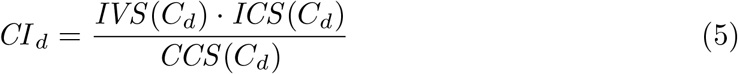

1. **Dense mode**: Among candidates meeting *τ*_in_, select the one with the smallest gain. This conservative strategy helps prevent sharp increases in *CI*_*d*_ in subsequent rounds.
2. **Standard mode**: Among candidates meeting *τ*_in_, select the one with the largest *w*_SSN_.

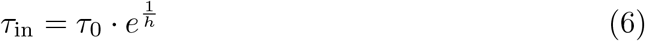

Add the selected node to *C*_*d*_, as illustrated in Figure 1b(v). If no candidate satisfies *τ*_in_, proceed directly to Step 5.

#### [Step 4] Pruning

Triggered periodically during community detection. TD-COM evaluates each node in *C*_*d*_: if its removal yields Δ*CI* ≥ *τ*_pr_, the node is flagged as a pruning candidate. The pruning candidate with maximum Δ*CI* is then pruned (Figure 1b(vi)).

#### [Step 5] Termination

The iteration terminates and outputs the final sample-specific DNB community *C*_*d*_. The DNBs within the resulting *C*_*d*_ constitutes the personalized EWS for the disease.

### 3.3 Determination of the critical state and its DNBs

Finally, we performed independent-samples *t*-tests to compare *CI*_*d*_ values between adjacent stages in disease progression. A significant increase in *CI* signals that the system has entered a state of heightened sensitivity to perturbations, which we term the critical state, a crucial time window for early intervention. For each sample in this critical state, the detected DNB community constitutes its single-sample DNB set. These sample-specific EWS enable precise interventions capable of halting or even reversing disease progression before it becomes irreversible. When DNBs appear frequently across critical-state samples, they serve as population-DNBs, representing shared EWS for the critical transition. Such population-DNBs are instrumental in defining critical-state subtypes and developing broad-spectrum intervention strategies.

## Acknowledgements

This work was supported in part by the National Natural Science Foundation of China (No. W2511070 (Fa Zhang), 32241027 (Fa Zhang), 62472034 (Xiaohua Wan), 62227807 (Bin Hu)), in part by the National Key R&D Program of China (No. 2019YFA0706200 (Bin Hu)).

